# An acylated N-terminus and a conserved loop regulate the activity of the ABHD17 de-acylase

**DOI:** 10.1101/2024.05.14.594217

**Authors:** Sydney Holme, Jennifer Sapia, Michael Davey, Stefano Vanni, Elizabeth Conibear

**Affiliations:** Department of Medical Genetics, University of British Columbia, Vancouver, BC VH6 3N1, Canada; Centre for Molecular Medicine and Therapeutics, British Columbia Children’s Hospital Research Institute, University of British Columbia, Vancouver, BC V5Z 4H4, Canada; Department of Biology, University of Fribourg, Chemin du Musée 10, CH-1700 Fribourg, Switzerland; Swiss National Center for Competence in Research (NCCR) Bio-inspired Materials, University of Fribourg, Chemin des Verdiers 4, CH-1700 Fribourg, Switzerland

## Abstract

The dynamic addition and removal of long chain fatty acids modulates protein function and localization. The alpha/beta hydrolase domain-containing (ABHD) 17 enzymes remove acyl chains from membrane localized proteins such as the oncoprotein NRas, but how the ABHD17 proteins are regulated is unknown. Here, we used cell-based studies and molecular dynamics simulations to show that ABHD17 activity is controlled by two mobile elements – an S-acylated N-terminal helix and a loop – that flank the substrate-binding pocket. S-acylation at multiple sites buries the helix in the membrane, which allows hydrophobic residues in the loop to interact with the bilayer. This stabilizes the conformation of both helix and loop, alters the conformation of the binding pocket and optimally positions the enzyme for substrate engagement. S-acylation may be a general feature of acyl-protein thioesterases. By providing a mechanistic understanding of how the lipid modification of a lipid-removing enzyme promotes its enzymatic activity, this work contributes to our understanding of cellular S-acylation cycles.

## Introduction

S-acylation, which involves the post-translational addition of a long chain fatty acid to cysteine residues, is the only reversible lipid modification. Dynamic S-acylation has been shown to influence protein localization, stability and activity and is important for cellular processes such as signaling and growth (reviewed in Chen *et al*., 2021 and S Mesquita *et al*., 2024). For example, NRas undergoes a cycle of acyl chain addition and removal that regulates its localization at the plasma membrane and is essential for oncogenic signaling (Remsberg *et al*., 2021; Xu *et al*., 2012). Altered acylation dynamics are implicated in conditions ranging from cancer and inflammation to neurodegenerative and cardiovascular diseases (Lin, 2021; Main & Fuller, 2022; Ramzan *et al*., 2023; Zhou *et al*., 2023).

Two classes of enzymes regulate S-acylation cycles: the protein-acyl transferase (PAT) enzymes that attach acyl chains to cytosolically-exposed cysteines via a thioester bond, and the acyl-protein thioesterase (APT) enzymes that remove the lipid (Greaves & Chamberlain, 2010; Won *et al*., 2018). While an estimated 10-20% of all human proteins are S-acylated, predominantly with the 16-carbon lipid palmitate (Muszbek *et al*., 1999), only a small subset of these undergoes rapid de-acylation (Martin *et al*., 2011). Cellular APT activity can also be enhanced by growth factor signaling (Kathayat *et al*., 2017), suggesting APT enzymes are both substrate-selective and stimulus-responsive. However, the mechanisms that regulate their substrate binding and activity are not well understood.

The best studied cytosolic APTs are members of the metabolic serine hydrolase family, including APT1, APT2 and the alpha/beta hydrolase domain-containing 17 proteins ABHD17A-C (collectively referred to here as ABHD17). ABHD17 is best characterized for activity on NRas and synaptic PSD-95 (Lin & Conibear, 2015; Yokoi *et al*., 2016), though the list of targets is growing (Dixon *et al*., 2023 *Preprint*; Remsberg *et al*., 2021; Ulengin-Talkish *et al*., 2021). Activity of ABHD17 on the stress-regulated exon variant of the large conductance voltage-and calcium-activated potassium (STREX BK) channel, nucleotide-binding domain, leucine-rich repeat, and pyrin domain-containing protein 3 (NRLP3), and microtubule-associated protein 6 (MAP6) implicates dynamic acylation in physiological processes such as cellular excitability, inflammatory disease and neuronal maturation and development (McClafferty *et al*., 2020; Tortosa *et al*., 2017; Zheng *et al*., 2023). Notably, a number of other enzymes have been proposed to have thioesterase activity and are under active investigation (Cao *et al*., 2019; Martin *et al*., 2011; Rosier *et al*., 2021; Savinainen *et al*., 2014; Xu *et al*., 2018; Yokoi *et al*., 2016).

From a structural and mechanistic point of view, an interesting feature of many APTs is that they themselves are acylated, which directs them to membranes where their substrates are located (Kong *et al*., 2013; Cao *et al*., 2019; Martin and Cravatt, 2009). Recent work suggests that acylation of APT2 has additional regulatory functions and occurs through a multistep process involving a membrane-targeting basic patch and hydrophobic beta-tongue (Abrami *et al*., 2021). These structures are required for subsequent acylation of an N-terminal cysteine that confers stable membrane binding and deforms the bilayer, facilitating substrate extraction.

It is not known if these are general principles that apply to other APTs, including the ABHD17 thioesterases. The acylated ABHD17 N-terminus is important for localization and activity (Lin & Conibear, 2015; Yokoi *et al*., 2016), but it remains unclear whether other regions of the protein are important for membrane targeting and substrate extraction. Here, we show that ABHD17A undergoes an activation process involving two distinct structural features. Recruitment of an N-terminal helix followed by acylation is essential for the insertion of a flexible and conserved loop into the bilayer. Membrane binding of the N-terminal helix and loop orients ABHD17A for optimal substrate insertion and activity.

## Results

### The ABHD17A N-terminus is sufficient for S-acylation and plasma membrane localization

Deletion of the ABHD17 N-terminus, or mutation of all 5 cysteines in this region, blocks S-acylation and plasma membrane targeting (Martin and Cravatt, 2009; Lin and Conibear, 2015; Yokoi *et al*., 2016). However, it is unclear if other regions of ABHD17 mediate the initial membrane binding that is required for acylation, as reported for APT2 (Abrami *et al*., 2021). To identify the minimal acylation determinant, we focused our studies on ABHD17A, which shows robust activity on NRas (Lin & Conibear, 2015), and used a sensitive bioluminescence resonance energy transfer (BRET) assay to quantitate its localization at different organelles (Lan *et al*., 2012). ABHD17A was tagged with the luciferase Rluc8 (donor) and expressed at low levels with different Venus-tagged organelle markers (acceptors). Proximity between the donor and acceptor constructs at organelle membranes results in a high net BRET ratio that reflects the subcellular localization of ABHD17A mutants. We found that replacing all five N-terminal cysteine residues with serine, or deleting the first 19 residues of ABHD17A, significantly decreased plasma membrane localization (Figure 1A), consistent with the results of previous immunofluorescence studies (Martin and Cravatt, 2009; Lin and Conibear, 2015). Both mutations also reduced localization of ABHD17A at the Golgi (Figure 1A).

**Figure 1.**
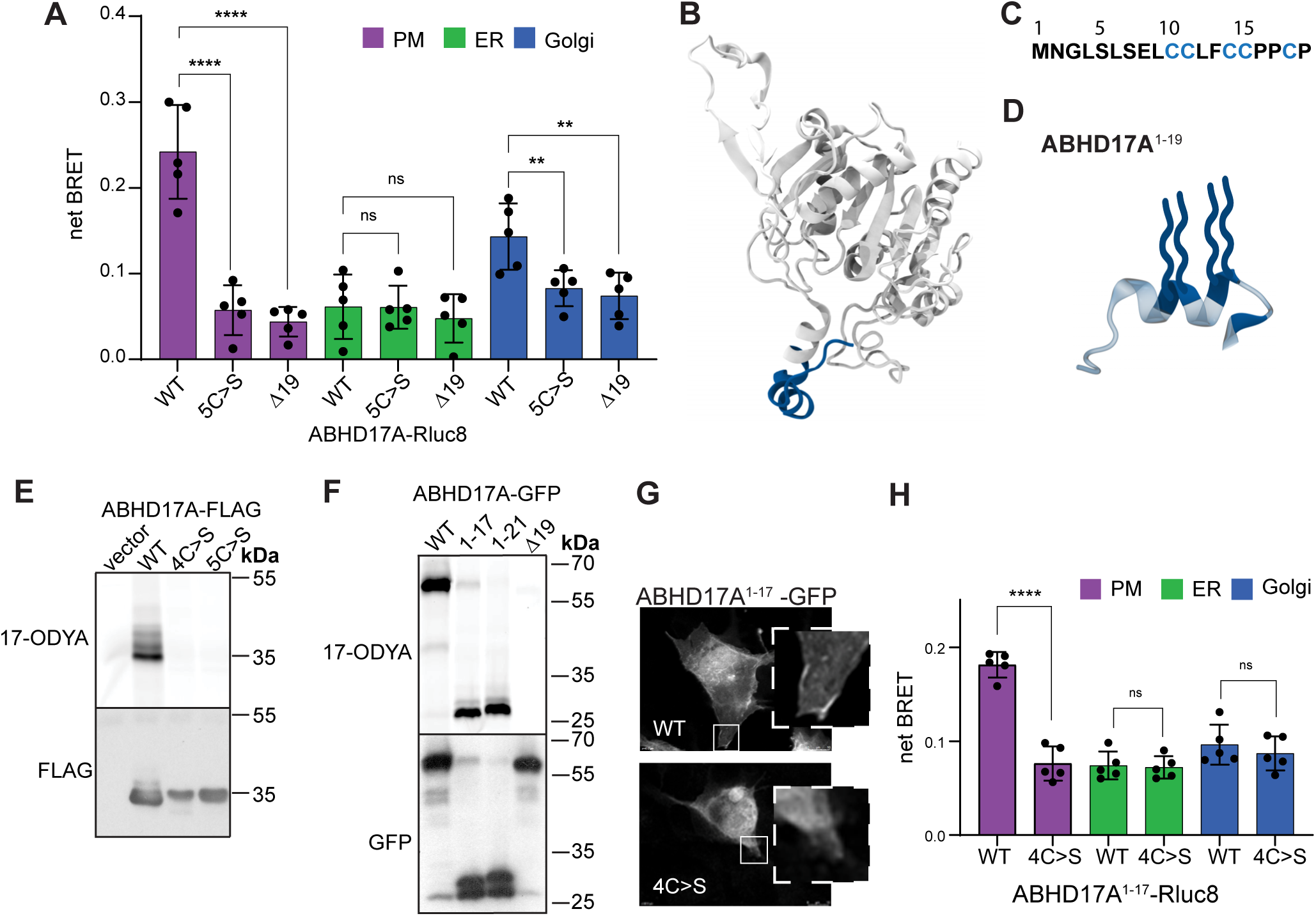
The N-terminus is necessary and sufficient for S-acylation and plasma membrane localization. **(A)** Plasma membrane localization of WT ABHD17A measured with BRET is lost with either mutation of N-terminal cysteine residues or deletion of the N-terminus. Cells were transiently transfected to co-express Rluc8-tagged ABHD17A mutants with a Venus-tagged organelle marker for the plasma membrane, Golgi or ER. One-way ANOVA with Tukey’s multiple comparison test; n=5, ns = p > 0.05, ** = p ≤ 0.01, **** = p ≤ 0.0001. Error bars depict standard deviation (STDEV). **(B)** AlphaFold2 (AF2) predicted structure of ABHD17A showing the first 19 residues in blue. **(C)** Sequence of the first 19 residues of ABHD17A showing the putative acylated cysteine residues in blue. **(D)** The ABHD17A N-terminus contains a predicted alpha-helix. Cysteine residues are shown in blue with acyl groups indicated by blue wavy lines attached to Cys10, 11, 14, 15. **(E)** Acylation of ABHD17A is abolished when either the first four or five N-terminal cysteine residues are mutated to serine as visualized with a click-labeling assay using the palmitate analog 17-octadecynoic acid (17-ODYA). Upper panel shows 17-ODYA signal and lower panel shows western blot of total protein detected with FLAG antibody. **(F)** The first 17 residues of ABHD17A (ABHD17A^1-17^) are sufficient for acylation. 17-ODYA signal was detected by click-labeling (upper panel); total protein was detected with GFP antibody (lower panel). **(G)** Plasma membrane localization of the first 17 residues of ABHD17A is lost when cysteine residues are mutated to serine. Representative confocal images of fixed cells are shown; the inset shows the plasma membrane localization of WT compared to the cytosolic signal of 4C>S. **(H)** Plasma membrane localization of the first 17 residues of ABHD17A requires N-terminal cysteine residues. The BRET assay described in *(A)*. One-way ANOVA with Tukey’s multiple comparison test; n=5, ns = p > 0.05, **** = p ≤ 0.0001. Error bars indicate STDEV.

AlphaFold2 (AF2) modeling (Jumper *et al*., 2021; Varadi *et al*., 2022) indicates that the N-terminal region of ABHD17A contains an alpha helix, with four of the five cysteine residues predicted to lie on or near one face of the helix (Figure 1B-D). By labeling cells with the ‘click-able’ palmitate analog 17-octadecynoic acid (17-ODYA), we found that mutation of these 4 cysteine residues was sufficient to abolish ABHD17A acylation (Figure 1E). Moreover, a truncated form of ABHD17A consisting of only its first 17 residues, which includes the 4 cysteines, was acylated as efficiently as a longer fragment that contains all 5 cysteine residues (Figure 1F). This minimal N-terminal domain (ABHD17A^1-17^ GFP) was detected at the plasma membrane both by confocal microscopy (Figure 1G) and by the BRET-based localization assay (Figure 1H), and its targeting was abolished by mutating all four cysteine residues to serine. Together, this demonstrates that the region containing the N-terminal helix is necessary and sufficient for ABHD17A S-acylation and plasma membrane localization, and that – in contrast to APT2 – regions outside of the acylation motif are not required.

### The hydrophobicity of the ABHD17A N-terminus is important for enzyme activity

The minimal N-terminal domain has 4 putative S-acylation sites, yet two membrane anchors are considered sufficient to confer stable membrane association to proteins (Shahinian and Silvius, 1995; Conibear & Davis, 2010). When mutated individually, no single cysteine residue was required for enzyme activity (Figure S1A, B). When we substituted two of the four cysteine residues at a time, the acylation of the resulting mutants was generally reduced to 50% of wild type levels, as predicted if all 4 cysteines are subject to modification (Figure S1C, D) and plasma membrane targeting, though reduced, was still observed (Figure S1F). However, specific combinations had strikingly different effects on enzyme activity: the C14,15S and C10,14S mutations largely blocked deacylation activity on NRas, whereas C10,11S and C11,15S mutations had relatively little impact (Figure 2A, B). This suggests that subtle differences in the position of the acyl-modified sites can have dramatic effects on activity (Figure S1G), and that the acylated N-terminus may be important for more than just membrane targeting.

**Figure 2.**
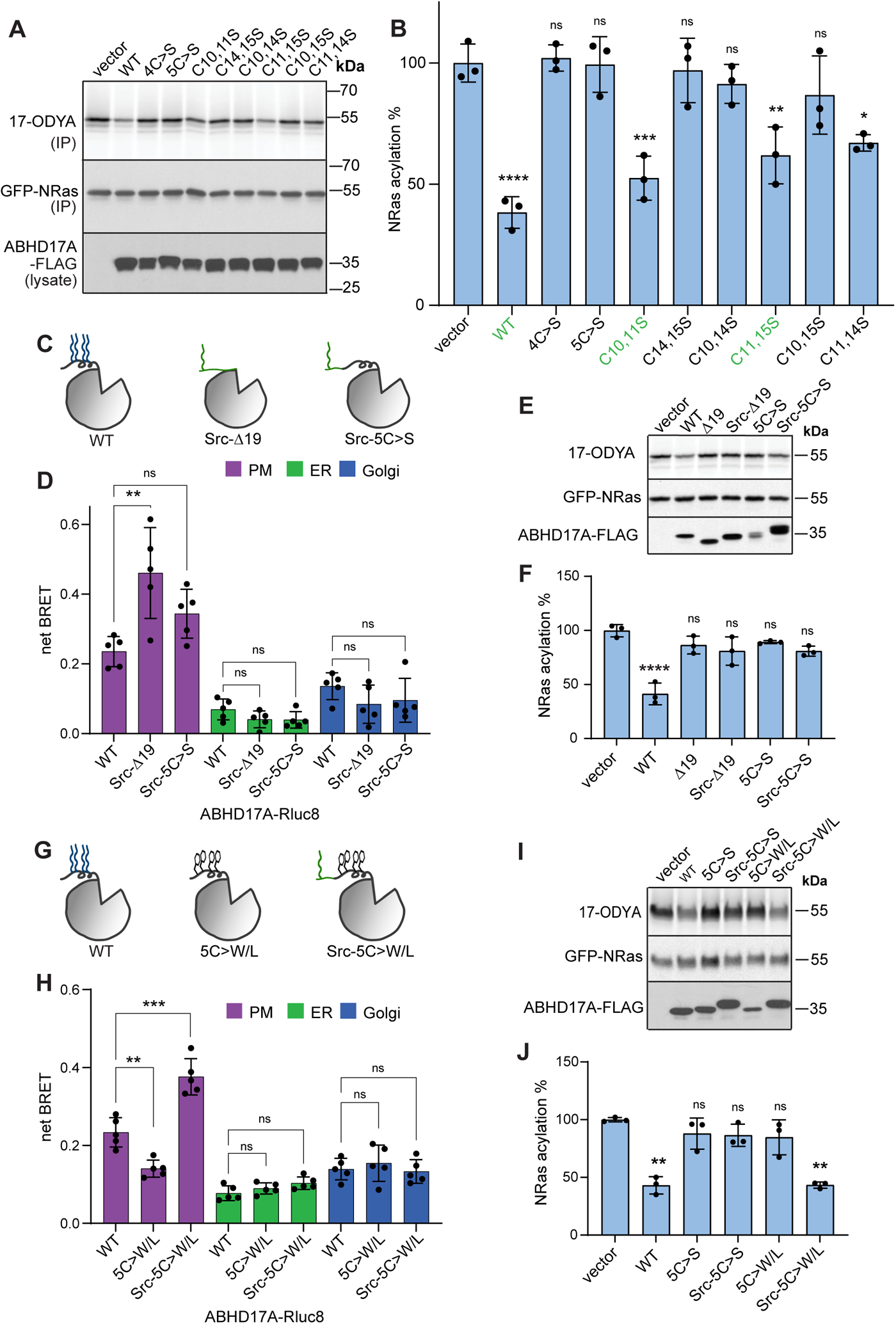
Hydrophobicity of the ABHD17A N-terminus is important for activity. **(A)** Specific cysteine combinations are required for the activity of FLAG-tagged ABHD17A on NRas. Acylation of GFP-NRas was detected by click-labeling with 17-ODYA (upper panel). Levels of immunoprecipitated NRas (IP; middle panel), or total ABHD17A (lysate; lower panel), were detected by western blot using anti GFP or FLAG antibody, respectively. **(B)** Quantification of GFP-NRas acylation shown in *(A)*. One-way ANOVA with Tukey’s multiple comparison test; n=3, ns = p > 0.05, * = p ≤ 0.05, ** = p ≤ 0.01. Statistical analysis shows comparison to vector. Error bars indicate STDEV. **(C)** Schematic of ABHD17A constructs. **(D)** The Src myristoylation motif restored plasma membrane localization of ABHD17A lacking either the first 19 residues, or N-terminal cysteine residues, as quantified by BRET analysis. One-way ANOVA with Tukey’s multiple comparison test; n=5, ns = p > 0.05, ** = p ≤ 0.01. Error bars indicate STDEV. **(E)** Recruiting non-acylated ABHD17A to the plasma membrane with a Src motif did not restore activity on NRas. Acylation of GFP-NRas was detected by click-labeling with 17-ODYA (upper panel). Levels of immunoprecipitated NRas (middle panel), or total ABHD17A (lower panel), were detected by western blot using anti GFP or FLAG antibody, respectively. **(F)** Quantification of GFP-NRas acylation in *(E).* One-way ANOVA with Tukey’s multiple comparison test; n=3, ns = p > 0.05, *** = p ≤ 0.001. Statistical analysis shows comparisons to vector. Error bars indicate STDEV. **(G)** Schematic of ABHD17A constructs with hydrophobic residues in place of N-terminal cysteine residues. **(H)** N-terminal hydrophobic residues localize ABHD17A to the plasma membrane when paired with the Src motif as quantified by BRET analysis. One-way ANOVA with Tukey’s multiple comparison test; n=5, ns = p > 0.05, ** = p ≤ 0.01, *** = p ≤ 0.001. Error bars indicate STDEV. **(I)** Attaching the Src motif to the ABHD17A mutant with hydrophobic N-terminal residues restored activity on GFP-NRas. Acylation of GFP-NRas was detected by click-labeling with 17-ODYA (upper panel). Levels of immunoprecipitated NRas (middle panel), or total ABHD17A (lower panel), were detected by western blot using anti GFP or FLAG antibody, respectively. **(J)** Quantification of NRas acylation in *(I).* One-way ANOVA with Tukey’s multiple comparison test; n=3, ns = p > 0.05, ** = p ≤ 0.01. Statistical analysis shows comparisons to vector. Error bars indicate STDEV.

Replacing the acylated N-terminus of ABHD17B with a plasma membrane anchor based on the myristoylated N-terminus of Src did not restore activity on the known substrate PSD-95, highlighting the functional importance of the ABHD17 N-terminus (Yokoi *et al*., 2016). We hypothesized that this heterologous plasma membrane targeting strategy might circumvent the need for acylation if the N-terminal sequences of ABHD17 were retained. Appending the Src myristoylation motif to either the full length, acyl-deficient version of ABHD17A (Src-5C>S-ABHD17; Figure 2C) or to the N-terminal deletion mutant (Src-Δ19-ABHD17) resulted in efficient plasma membrane targeting in a BRET assay (Figure 2D). However, neither construct showed significant deacylation activity on NRas (Figure 2E, F).

Because ABHD17A activity is sensitive to the position of acylation sites along the N-terminal helix (Figure S1G), we speculated that acylation creates a hydrophobic face that is important for the correct positioning of the helix at the membrane. To test this idea, we created a version of ABHD17A that has bulky hydrophobic residues (Trp/Leu) in place of the acylated cysteines (Figure 2G). When the Src motif was attached to the N-terminus (Src-5C>W/L-ABHD17), this mutant was localized to the plasma membrane (Figure 2H) and its activity on NRas, as measured by click assay, was significantly restored (Figure 2I, J). This suggests the hydrophobicity of the N-terminal helix is essential for ABHD17A activity.

### ABHD17A interacts with membranes with two distinct domains: the N-terminal helix and a conserved loop

Our results indicate that the thioesterase activity of ABHD17A depends on a specific interaction between its N-terminal helix (“N-helix”) and the lipid bilayer. To understand how ABHD17A interacts with membranes, we turned to molecular dynamics (MD) simulations. Coarse-Grained (CG) simulations using the MARTINI force field (Souza *et al*., 2021) were initially employed to investigate the interaction between the predicted AF2 model of ABHD17A and a PM-like model membrane, as this methodology has been shown to accurately predict protein-membrane interfaces for peripheral proteins (Srinivasan *et al*., 2021; Srinivasan *et al*., 2023 *Preprint*) and it has been successfully used to investigate the membrane-binding interface of the thioesterase APT2 (Abrami *et al*., 2021). We observed that the enzyme primarily interacts with the membrane through two regions: the N-helix and a structurally adjacent loop (residues 222-233, Figure 3A), that is conserved across metazoans (Fig 3B). Quantitative analysis of protein-membrane contacts and protein occupancy are shown in Figures 3C and D.

**Figure 3.**
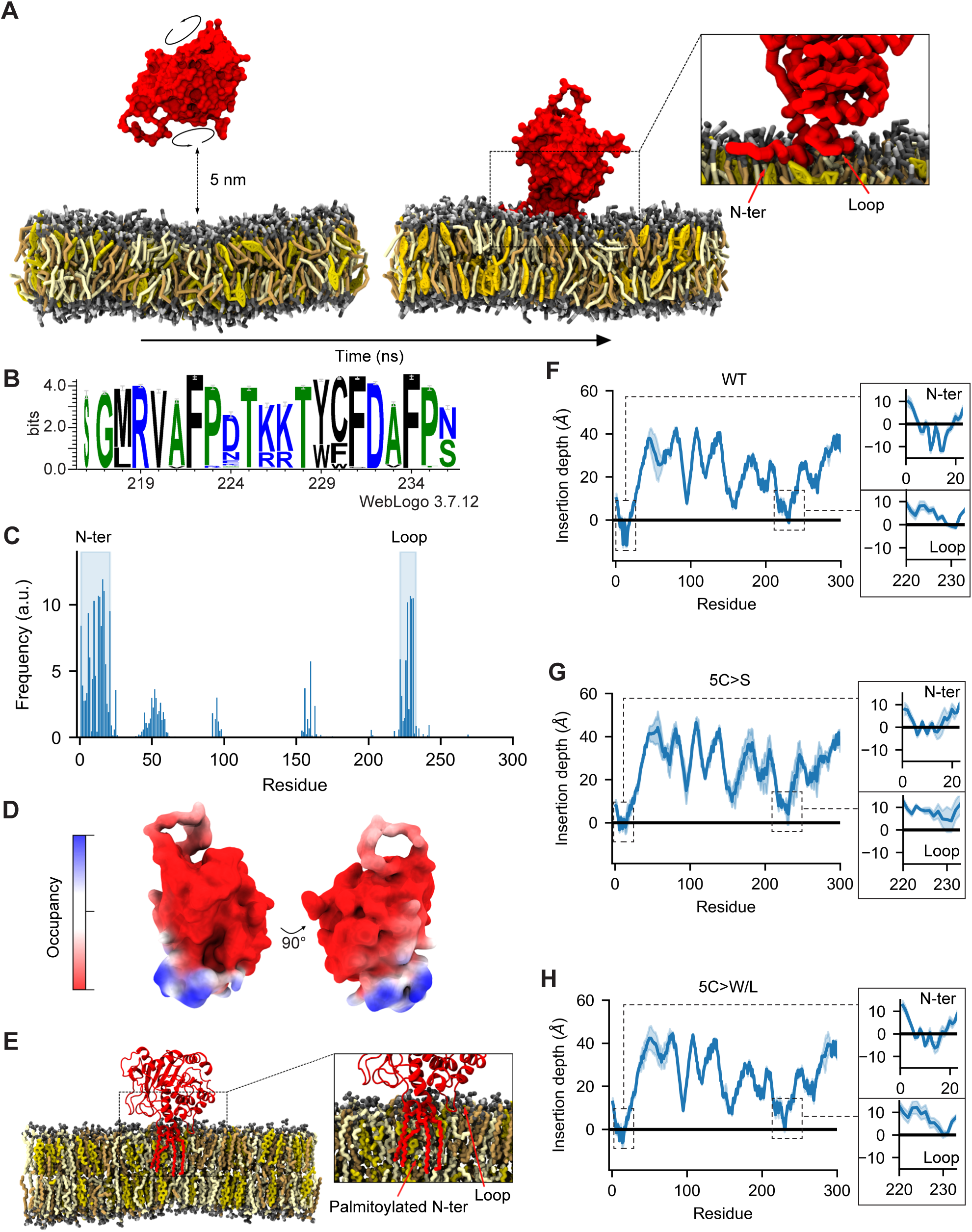
ABHD17A-membrane interaction occurs through two distinct domains. **(A)** Representative mechanism for the binding of ABHD17A (red) to a PM-like membrane (POPC: light yellow, POPS: orange, CHOL: yellow) in CG simulations: the protein, initially randomly oriented at a minimum distance of 5 nm from the bilayer, interacts with the membrane via both its N-terminal helix and the adjacent loop. **(B)** Conservation of the loop residues in the ABHD17 proteins in metazoans as shown with Weblogo (Crooks *et al*., 2004). **(C-D)** Membrane interaction of the N-terminal helix and conserved loop shown in the contacts analysis (C) and the occupancy maps (D)**. (E)** Snapshot from a representative AA simulation, showing the back-mapped atomistic structure of ABHD17A with the addition of palmitoyl-tails to cysteine residues at its N-terminus. (**F)** Insertion depth analysis of AA simulations confirms that ABHD17A inserts into the membrane via both the palmitoylated-N-terminus (residues 1-21) and the adjacent loop (residues 222 to 233). **(G)** Insertion depth analysis of ABHD17A 5C>S with AA simulations shows mutation of the five N-terminal cysteine residues to serine decreases both N-terminal and loop membrane insertion. **(H)** Restoring N-terminal hydrophobicity of the non-acylated ABHD17A (5C>W/L) rescues insertion depth of the N-terminus and loop in AA simulations.

To gain deeper insights into ABHD17A’s interaction with the PM-like membrane model, we next opted to adopt a finer resolution by means of MD simulations at the atomistic level. To this end, a representative snapshot from the CG simulations was back-mapped to all-atom (AA) and later palmitoylated at four cysteine sites (Figure 3E). AA-MD simulations of this configuration showed that the CG binding-pose remained almost intact, with both the N-terminus (first 21 residues) and the adjacent loop (residues 222-233) penetrating the membrane to an average depth of approximately 11 Å and 1 Å, respectively, with respect to the average level of the phosphate groups of the monolayer phospholipids (Figure 3F).

Our simulations predict that both N-helix and loop are important for the interaction of ABDH17A with the membrane. To understand the relative contribution of the two regions, we used AA-MD simulations to model different ABHD17A mutants. Upon mutation of the N-terminal cysteine residues to serine (5C>S), which renders the protein inactive (Figure 2E, F), the binding mode of ABHD17 was altered: the insertion depth of the N-terminus decreased significantly (to about 2 Å) and the loop was no longer inserted into the membrane (Figure 3G). In agreement with the functional assays (Figure 2I, J), mutation of the N-terminal cysteines to bulkier and more hydrophobic residues such as tryptophan (5C>W/L) partially restored the membrane insertion of both the N-helix (to about 5Å) and loop (to about 0.2 Å; Figure 3H). Overall, our data suggest that membrane insertion of the palmitoylated N-helix allows the adjacent loop to insert into the bilayer, and that both interactions contribute to enzyme activity.

### Hydrophobic loop residues are important for ABHD17A activity

To test the importance of the conserved loop structure observed in the MD simulations for deacylation activity, we created a series of mutants with four consecutive alanine substitutions spanning the loop (Figure 4A). Mutations in the central portion of the loop, which contains polar and charged residues, did not affect enzyme activity on NRas, but mutating outer regions of the loop abolished activity (Figure 4B, C). These outer regions contain three conserved hydrophobic residues (F222, Y229, F231) that lie in a part of the loop that is predicted to insert into the membrane (Figure 3A) with protruding side chains that could contribute to membrane binding (Figure 4D).

**Figure 4.**
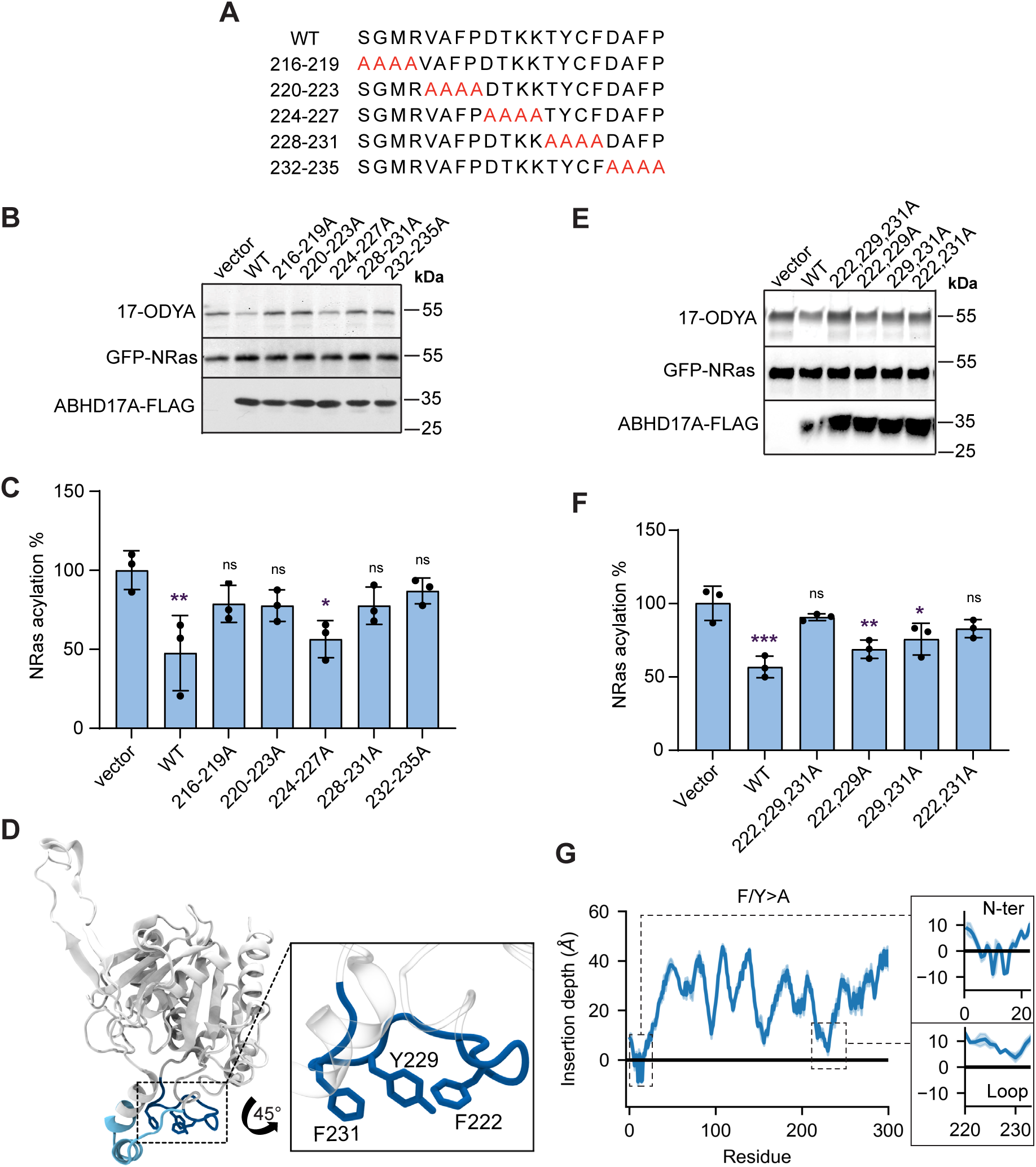
Hydrophobic loop residues are important for activity. **(A)** Schematic of the alanine scan used to determine the regions of the loop required for ABDH17A activity. (B) Alanine loop mutants show flanking regions of the loop are important for ABHD17A activity on NRas. Activity of ABHD17A was measured by acylation of GFP-NRas as detected by click-labeling with 17-ODYA (upper panel). Levels of immunoprecipitated NRas (middle panel), or total ABHD17A (lower panel), were detected by western blot using anti GFP or FLAG antibody, respectively. **(C)** Quantification of GFP-NRas acylation in *(B)*. One-way ANOVA with Tukey’s multiple comparison test; n=3, ns = p > 0.05, * = p ≤ 0.05. Statistical analysis depicts comparison to vector. Error bars indicate STDEV. **(D)** AF2 predicted structure of ABHD17A showing the loop adjacent to the active site in blue. Inset shows a close up of the loop and the orientation of the side chains of the three hydrophobic loop residues F222, Y229 and F231. **(E)** Three hydrophobic loop residues (F222, Y229, F231) are needed for ABHD17A activity on GFP-NRas. Acylation of GFP-NRas was detected by click-labeling with 17-ODYA (upper panel). Levels of immunoprecipitated NRas (middle panel), or total ABHD17A (lower panel), were detected by western blot using anti GFP or FLAG antibody, respectively. **(F)** Quantification of GFP-NRas acylation in *(E)*. One-way ANOVA with Tukey’s multiple comparison test; n=3, ns = p > 0.05, * = p ≤ 0.05, ** = p ≤ 0.01. Statistical analysis depicts comparison to vector. Error bars indicate STDEV. **(G)** Insertion depth analysis shows that mutating hydrophobic loop residues 222, 229, 231 to alanine in AA simulations of acylated ABHD17A impairs membrane insertion of the conserved loop.

To determine if these hydrophobic loop residues are required for activity, we further made alanine mutations in different combinations (Figure 4E). Most of the single and double mutations had relatively minor effects, with F222 and F231 having the greatest effect when mutated together. However, mutating all three hydrophobic residues abolished activity (Figure 4F). In AA-MD simulations using the palmitoylated form of ABHD17A (Figure 3E), mutation of these hydrophobic loop residues (F/Y>A) resulted in a total loss of membrane insertion for the loop, without affecting binding of the N-terminus (Figure 4G). Taken together, these results suggest that the integrity of both the N-helix and loop are important for membrane binding and activity of ABHD17A.

### Mutation of the N-terminal helix and conserved loop alter binding pocket conformation

Many metabolic serine hydrolases have a lid domain that regulates access to the substrate-binding pocket (Devedjiev *et al*., 2000; Filippova *et al*., 2013). However, the substrate-binding site of ABHD17, and the mechanisms that regulate substrate binding, are not known. Using a binding pocket prediction algorithm relying on Voronoi tessellation, called FPocket (Le Guilloux *et al*., 2009), we identified a hydrophobic cavity that represents a potential substrate-binding site (Figure 5A). This cavity appears as a cleft in the AF2-predicted structure (Figure 5B), with the opening next to the membrane, and the catalytic serine located within the part of the cleft that is distal to the membrane. Based on our top-scoring SwissDock model (Figure 5B-D; Grosdidier *et al*., 2011a), palmitate is predicted to dock in this cleft, with its carboxylate group adjacent to the catalytic serine (Ser190), as would be expected for substrate depalmitoylation. The residues from the N-helix and loop frame this binding pocket opening, creating what appears to be a channel for substrate entry (Figure 5D). Through MD simulations, we determined that the N-helix and loop regions of ABHD17A are flexible regions that can adopt a wide range of conformations in solution (Figure S2A, B), but upon membrane association they appear to lock into a preferred conformation that could favor substrate insertion into the hydrophobic pocket (Figure S2B-E). Thus, dynamic changes in the positions of N-helix and/or loop have the potential to control access to the substrate binding site.

**Figure 5.**
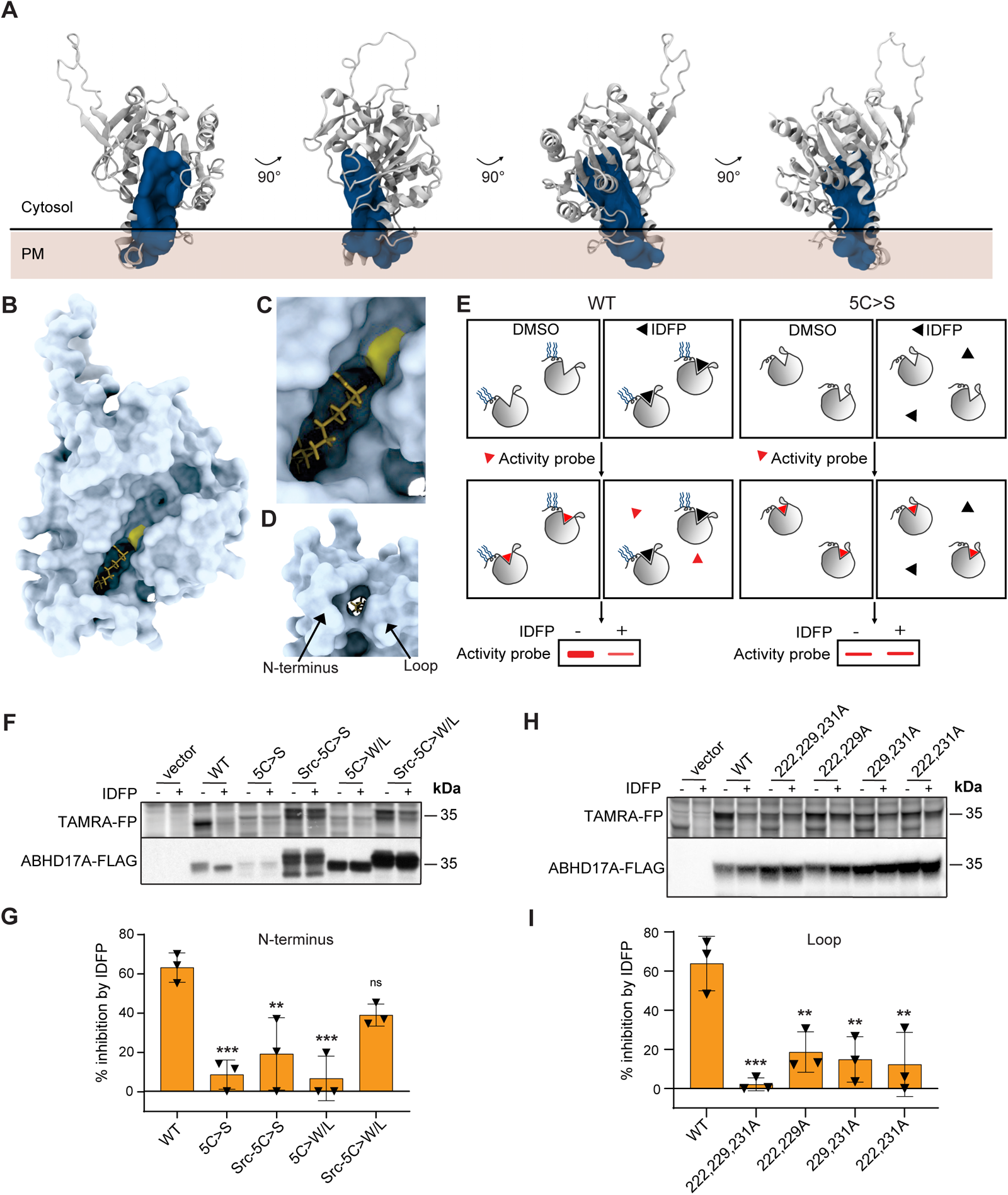
The N-terminus and loop affect binding pocket conformation. **(A)** Pocket analysis identifies one major cavity inside ABHD17A, indicated by the blue solid surface. Its localization close to the membrane interface suggests that the cavity is a potential substrate binding pocket for ABHD17A. **(B-D)** An ABHD17A surface model of the AF2 predicted structure reveals a prominent cleft. Palmitic acid is docked into this cleft by SwissDock (Grosdidier *et al*, 2011). The active site serine is shown in yellow. **(C)** Close up of the top-scoring SwissDock model showing palmitic acid interacting with ABHD17A. **(D)** View of the membrane-contacting surface shows the N-terminus and loop are predicted to frame the entrance of the channel. **(E)** Schematic of the cABPP assay used to determine substrate binding of ABHD17A mutants. Cell lysates expressing ABHD17A-FLAG constructs are incubated with the 12-carbon fluorophosphonate inhibitor IDFP or dimethyl sulfoxide (DMSO). Subsequent incubation with the fluorescent activity probe TAMRA-FP covalently labels the catalytic serine of ABHD17A molecules that have not bound the inhibitor. IDFP binding causes a reduction of in-gel fluorescence (left side). Mutations that alter the conformation of the substrate-binding pocket reduce IDFP binding, thus no change in fluorescence is seen (right side). **(F)** The Src myristoylation motif paired with hydrophobic N-terminal residues partially restored the binding pocket conformation of non-acylated ABHD17A mutants. TAMRA-FP binding was visualized with in-gel fluorescence (upper panel) and total protein was detected by western blot using an anti-FLAG antibody (lower panel). **(G)** Quantification of *(F)* depicted by percentage of IDFP inhibition. One-way ANOVA with Tukey’s multiple comparison test; n=3, ns = p > 0.05, ** = p ≤ 0.01, *** = p ≤ 0.001. Statistical analysis shows comparison to wildtype ABHD17A. Error bars indicate STDEV. **(H)** Mutation of three hydrophobic loop residues (F222, Y229, F231) significantly decreased IDFP binding by cABPP analysis. **(I)** Quantification of *(H)* depicted by percentage of IDFP inhibition. One-way ANOVA with Tukey’s multiple comparison test; n=3, ** = p ≤ 0.01, *** = p ≤ 0.001. Statistical analysis shows comparison to wildtype ABHD17A. Error bars indicate STDEV.

Similar to other mSHs, acyl protein thioesterases are irreversibly inhibited by fluorophosphonate (FP) probes, which form a covalent linkage with the active site serine (Creighton, 1993; Piñeiro-Sánchez *et al*., 1997; Blankman *et al*., 2007; Martin *et al*., 2011). To examine the role of the N-helix and loop in regulating substrate binding, we used a competitive activity-based protein profiling (cABPP) assay that measures the ability of a pseudosubstrate FP inhibitor to bind at the active site and prevent subsequent labeling by the fluorescent FP probe TAMRA-FP (Leung *et al*., 2003; Figure 5E). We found that preincubation with the 12-carbon inhibitor isopropyl dodecylfluorophosphonate (IDFP) effectively inhibited TAMRA-FP binding to wild type ABHD17A, but not to inactive forms of ABHD17A, such as 5C>S, or the plasma membrane targeted form of this mutant (Src-5C>S-ABHD17A; Figure 5F-I). The altered substrate binding properties of these mutants suggest that changes to either the N-helix or loop that affect their membrane insertion alters the accessibility or conformation of the substrate binding pocket.

We hypothesized that restoring the hydrophobicity of the N-helix by substituting cysteines for bulky hydrophobic residues, which significantly enhanced activity (Figure 2H), would also restore substrate binding. Indeed, this mutant (Src-5C>W/L-ABHD17A) showed significantly restored IDFP inhibition (Figure 5F, G), suggesting that the insertion of the N-helix in the bilayer is critical for binding pocket accessibility. Moreover, mutation of all three hydrophobic loop residues (F222, Y229, F231), which blocked ABDH17A activity, significantly decreased IDFP binding (Figure 5H, I). This was not observed when hydrophobic loop residues were mutated individually, supporting the idea that their contribution is partially redundant (Figure S2F-I).

Taken together, these results show that mutations that reduce the membrane insertion of either the N-helix or loop alter the conformation of the substrate binding pocket and impair deacylation activity. Conversely, compensatory changes that restore membrane insertion restore pocket conformation and enzyme activity. This supports the model that ABHD17A uses both the N-terminal helix and a conserved loop to bind membranes and orient the hydrophobic pocket in a way that promotes substrate binding.

## Discussion

We have shown that the activity of ABHD17A relies on the membrane embedding of two regions that flank the substrate binding pocket: the acylated N-terminal helix and a flexible loop. Both elements must be inserted into the membrane to position the substrate binding pocket at the bilayer interface in a way that facilitates substrate entry. While our analysis focused on ABHD17A, all three ABHD17 proteins are predicted to have a similar structure that includes an acylated N-terminus and conserved loop region and are likely to share the same regulatory mechanism.

### Acylation and membrane binding of the N-terminal helix

To be acylated, APTs must first encounter membrane-localized palmitoyltransferases (DHHC-PATs). In the case of APT2, two different regions – a basic patch and a hydrophobic ‘tongue’ – make this initial membrane contact, which leads to acylation of a single site near the N-terminus (Abrami *et al*., 2021). In contrast, our work suggests that acylation of the ABHD17 N-terminal domain, which contains a cluster of cysteines on or near a predicted alpha helix, does not require other domains. The amphipathic nature of this helix, together with the sulfhydryl side chains of cysteine residues, could favour insertion into the lipid core of the bilayer (Iyer & Mahalakshmi, 2019) and promote the initial membrane interaction needed for ABHD17 to encounter DHHC-PATs.

Because acylation is critical for membrane localization, the dynamic addition and removal of the acyl group was proposed to regulate the activity of APTs by altering access to membrane-localized substrates (Vartak *et al*., 2014), limiting their concentration by auto-deacylation in trans (Kong *et al*., 2013), or exposing lysine residues that target them for ubiquitination and degradation (Abrami *et al*., 2021). However, the acylation of ABHD17 on a cluster of 4 or 5 cysteines is predicted to shield it from APTs (Anwar & van der Goot, 2023), which would have to cleave multiple acyl groups to effect membrane release. Thus, ABHD17 is unlikely to be regulated by rapid cycling on and off membranes. Instead, we found that acyl chain positioning on the helix was important for activity. Although the reason for this was unclear, the spacing of acyl groups could influence helix orientation, subcellular localization or membrane domain association (Batrouni *et al*., 2021; El-Husseini *et al*., 2001).

While the acylated N-terminal helix of ABHD17 is important for membrane targeting, replacing it with a heterologous localization signal compromised activity (Yokoi *et al*., 2016), suggesting it has another role. We found that full deacylation activity requires the N-terminal helix to associate with the membrane through hydrophobic interactions. In molecular dynamics simulations, the acylated N-terminal helix inserts deeply into the bilayer, which in turn promotes the membrane insertion of the adjacent loop structure. While the insertion of the N-terminal helix is important for loop insertion, the converse is not true: mutations in the loop that block membrane binding and activity do not disrupt insertion of the N-helix. This suggests that acylation of the N-helix is an initial step that positions ABHD17 for loop insertion and substrate engagement.

The opening of the binding pocket faces the membrane, whereas the catalytic serine is at the base of the pocket, suggesting the lipid substrate must be extracted from the bilayer to enter the binding site. We hypothesize that the membrane insertion of both the N-helix and the loop, which flank the binding pocket, are important for orienting the enzyme so that the binding pocket is optimally positioned for lipid extraction to occur. Furthermore, insertion of the acylated N-helix is expected to perturb the bilayer, facilitating this extraction. Indeed, palmitate, the predominant lipid used for acylation, decreases membrane fluidity and destabilizes membranes in liposomes (Leekumjorn *et al*., 2009). Because both tryptophan and phenylalanine insertion also destabilize membranes (Popova *et al*., 2002), loop binding could also contribute to substrate extraction. Similarly, membrane deformation by APT2 is proposed to facilitate substrate removal and subsequent hydrolysis. (Abrami *et al*., 2021). Thus, acylation may be a common feature of APTs that not only serves as a membrane targeting signal, but also contributes to deacylation activity.

### Binding pocket conformation is regulated at the membrane

It is unclear if access to the ABHD17 pocket is dynamically regulated. Many lipases have highly dynamic lid structures that occlude the binding pocket until interaction with a membrane causes the lid to open (Khan *et al*., 2017). Lid opening and substrate engagement can also cause conformational changes in the binding pocket that promote binding and position the active site residues for cleavage. The APT1/2-like protein FTT258 found in the bacteria *Francisella tularensis* has a flexible loop that acts like a lid to regulate membrane binding, binding pocket accessibility and enzyme activity (Filippova *et al*., 2013; Smith *et al*., 2018). APT1 lacks a lid-like structure but does have a loop that forms part of the binding pocket and undergoes subtle structural rearrangements on substrate binding to accommodate the substrate, which provide isoform selectivity (Won *et al*., 2016; Harris *et al*., 2024). APT homologs in plants are also suggested to use sequences adjacent to their active site to select the chain length of substrates (Bürger *et al*., 2017).

Although ABHD17 lacks an obvious lid structure, we found the integrity of both the N-terminus and loop to be important for the conformation of the binding pocket. In MD simulations, the relative position of the N-helix and loop is highly variable in solution, but uniform when bound to membranes. Based on these observations, we favor the hypothesis that the binding and insertion of the N-helix and loop elements into the lipid bilayer stabilizes a conformation that promotes substrate engagement. The location of the active site serine in a deep cleft might help accommodate the acylated peptide as it slides into the active site. Whether ABHD17 proteins undergo additional structural rearrangements in response to substrate binding will be the topic of future studies.

Different organelle membranes have distinct lipid compositions and lipid packing properties that can influence protein binding and insertion (Bigay & Antonny, 2012). It is tempting to speculate that the need for correct insertion of the N-terminal helix and loop regions limits ABHD17 activity to specific organelles or membrane subdomains. This could help explain why ABHD17 appears to act primarily on substrates at the plasma membrane (Remsberg *et al*., 2021). How ABHD17 localization affects activity and substrate specificity, and whether ABHD17 proteins are also regulated by post-translational modification or expression levels (Abazari *et al*., 2023; Wild *et al*., 2023) are important areas of future research. A complete understanding of ABHD17 regulation will be helpful in developing therapeutic approaches that target ABHD17 and will increase our understanding of related thioesterases.

## Materials and Methods

### Plasmids

Venus-tagged KRas-tail, PTP1b-tail, Giantin, and Rluc8-N1 plasmids were gifts from Dr. Nevin Lambert (Augusta University, Georgia; Lan *et al*., 2012). ABHD17A mutant plasmids (plasmids 5-21 in Supplementary Table 1) were made with NEBuilder HiFi DNA Assembly (New England Biolabs), using primers (Supplementary Table 2) to amplify and mutate ABHD17A sequences that were inserted into EcoRI/Bsu36I-digested pABHD17A-FLAG (Lin & Conibear, 2015). The loop mutant plasmids (plasmids 22-33) were made by ligating two PCR products with SacII/BmgBI digested pABHD17A-FLAG. A similar approach was used to create pSrc-Δ19-FLAG, ligating the PCR product into EcoRI/XhoI digested pABHD17A(Δ19)-FLAG.

To create pABHD17A-EGFP, EGFP PCR product was ligated into EcoRi/SalI digested pABHD17A-FLAG. NEBuilder HiFi DNA Assembly was used to ligate 1-17 and 1-21 ABHD17A fragments with pABHD17A-EGFP that was digested with EcoRI/AgeI to remove ABHD17 sequences. pABHD17A-Rluc8 constructs were created by PCR amplifying ABHD17A mutants described above. Ligation of the PCR product with EcoRI/BamHI digested pRluc8-N1 was performed with NEBuider HiFi DNA Assembly.

### Cell culture and cDNA transfection

COS-7 cells from ATCC were maintained and propagated in high-glucose Dulbecco’s Modified Eagle Medium (DMEM; Gibco) supplemented with 10% fetal bovine serum (FBS; Gibco), 4 mM L-glutamine and 1 mM sodium pyruvate, in a humidified incubator at 37°C, 5% CO2.

COS-7 cells were grown on 60mm plates and transfected at 90% confluence using Lipofectamine 2000 as per manufacturer’s instructions with 1μg cDNA per plate of each DNA construct for 17-ODYA and cABPP labeling studies. For BRET, cells were grown in 6-well plates and transfected at 90% confluence with 0.5μg cDNA of acceptor constructs and 0.05μg cDNA of donor constructs. For fluorescence microscopy, cells were grown on 12mm glass coverslips (Thermo Fisher) in 24-well plates and transfected at 90% confluence with 0.4μg cDNA.

### BRET assays

Cells were washed in PBS, harvested with 0.25% EDTA-Trypsin (Gibco), resuspended in HBSS (Gibco) and transferred to opaque 96-well plates. Fluorescence and luminescence measurements were made using a Tecan Spark multimode microplate reader (Tecan). Fluorescence emission was measured at 485nm and 530nm. Coelenterazine h (5μM; Cayman) was added to cells immediately prior to making measurements. Raw BRET signals were calculated as the emission intensity at 530nm divided by the emission intensity at 485nm. Net BRET is the emission_acceptor_/emission_donor_ ratio minus the emission_acceptor_/emission_donor_ ratio measured from cells expressing only the BRET donor.

### 17-ODYA metabolic labeling

20 hours following transfection, COS-7 cells were washed once in PBS and starved for one hour in Base Labelling Media (1mM L-Glutamine, 1mM cysteine and 1mM sodium pyruvate in cysteine- and methionine-free DMEM). A 30μM solution of saponified 17-Octadecynoic acid (17-ODYA; Cayman) was prepared by first incubating 6mM ODYA in 7.2 mM sodium hydroxide at 65°C for 15 minutes, then adding this to Base Labeling Media containing 20% fatty acid-free BSA. After two hours in the 17-ODYA solution, cells were washed three times in PBS and placed at - 80°C for 24 hours. Cells were lysed with triethanolamine (TEA) lysis buffer [1% TX-100, 150 mM NaCl, 50 mM TEA pH 7.4, 2×EDTA-free Halt Protease Inhibitor (Life Technologies)] and vigorously mixed by pipetting. Lysates were cleared by centrifugation at 16,000× g for 15 min at 4°C, and the supernatant protein content was quantified using Bicinchoninic acid (BCA) assay (Life Technologies). 100 – 130 μg of supernatant was diluted in SDS-sample buffer (8% SDS, 4% Bromophenol Blue, 200 mM Tris-HCl pH 6.8, 40% Glycerol, 1% 2-mercaptoethanol). Samples were heated for 5 min at 95°C and kept at -20°C. The remaining lysate was used for immunoprecipitation and sequential on-bead CuAAC/click chemistry.

### Immunoprecipitation and sequential on-bead CuAAC/click chemistry

For immunoprecipitations, Protein A Sepharose beads (GE Healthcare) were washed three times in TEA lysis buffer and pre-incubated with rabbit anti-GFP antibody for 4 hours at 4°C, before the addition 500 μg – 1 mg of transfected COS-7 cell lysates. Immunoprecipitations were carried out for 12–16 hr on an end-to-end rotator at 4°C. Sepharose beads were then washed three times in modified RIPA buffer [150 mM NaCl, 1% sodium deoxycholate (w/v), 1% TX-100, 0.1% SDS, 50 mM TEA pH7.4].

Sequential on-bead click chemistry of immunoprecipitated 17-ODYA-labeled proteins was carried out as previously described (Zhang *et al*., 2010), with minor modifications. Sepharose beads were washed three times in RIPA buffer, and on-bead conjugation of AF647-azide (Invitrogen) to 17-ODYA was carried out for 1 hr at room temperature in 50 μL of freshly mixed click chemistry reaction mixture containing 1 mM TCEP (Aldrich), 1 mM CuSO^4^·5H_2_O (Sigma), 100 μM TBTA (Aldrich), and 100 μM AF647-azide in water. Beads were washed three times with RIPA buffer and resuspended in SDS buffer (150 mM NaCl, 12% SDS, 50 mM TEA pH7.4, 4% Bromophenol Blue, 200 mM Tris-HCl pH 6.8, 40% Glycerol and 1% 2-mercaptoethanol). Samples were heated for 5 min at 95°C and separated on 10% tris-glycine SDS-PAGE gels for subsequent in-gel fluorescence analyses, then transferred onto nitrocellulose membrane for Western blotting. The percent of acylated NRas was calculated using the ratio of ODYA to total purified NRas, normalized to the average value of the vector control.

Lysate samples frozen in SDS-sample buffer were thawed, heated at 95°C for two min, sonicated gently, and run on 13% tris-glycine SDS-PAGE gels for western blot analyses.

### Competitive activity-base protein profiling

24 hours following transfection with ABHD17A constructs, COS-7 cells were washed once in PBS, incubated with 0.25% trypsin for 3 min at 37°C, detached with DMEM + FBS (10%) and collected by centrifugation at 1000 g for 3 min. The cells were resuspended in 300μL 50mM Tris and lysed with gentle sonication on ice. Protein was quantified by BCA assay and 30μg protein was incubated either with DMSO or isopropyl dodecylfluorophosphonate (IDFP, 20 μM) at room temperature for 30 min. TAMRA-FP (2μM final concentration) labeling was carried out at room temperature for 1 hr and quenched with 4× SDS-sample buffer heated to 95°C for 5 min. Samples were separated on SDS–PAGE, analyzed by in-gel fluorescence, then transferred onto nitrocellulose membrane for Western blotting. Inhibition was calculated by dividing the TAMRA-FP signal treated with IDFP by the DMSO control and subtracting this from 100.

### In-gel fluorescence analyses

A Typhoon Trio scanner (GE Healthcare) was used to measure in-gel fluorescence of SDS– PAGE gels. AF647 signals were acquired using the red laser (excitation 633 nm) with a 670BP30 emission filter, and rhodamine signals were acquired with the green laser (excitation 532 nm), with a 580BP30 emission filter. Signals were acquired in the linear range and quantified using Fiji (Schindelin *et al*., 2012).

### Western blotting

Nitrocellulose membranes (50-206-3328; Fisher Scientific) were blocked in PBST [PBS with 0.1% Tween-20 (Sigma)] containing 3% bovine serum albumin (BSA, Sigma) for 1 hr. Membranes were then blotted with corresponding primary antibodies [mouse anti-GFP (11814460001; Roche) or rabbit anti-FLAG (701629; Thermo Fisher)] in PBST + 3% BSA for 1 hr followed by either horseradish peroxidase-conjugated polyclonal goat anti-mouse (115–035-146; Jackson ImmunoResearch Laboratories) or horseradish peroxidase-conjugated polyclonal goat anti-rabbit (170-6515; Biorad) in PBST + 3% BSA for 1 hr. All blots were developed with ECL chemiluminescent reagents (RPN2106; Cytiva) and exposed to Amersham Hyperfilm (CA95-17; VWR). Developed films were scanned, and densitometry performed in Fiji.

### Weblogo

ABHD17A homologs were identified at the metazoan level with OrthoDB (Kuznetsov *et al*., 2022) and aligned with MUSCLE (Edgar, 2004). Weblogo (Crooks *et al*., 2004) was used to analyze amino acids 216-235 of *H. sapiens* ABHD17A.

### Fluorescence microscopy image acquisition and processing

Coverslips were mounted onto a microscopy slide using ProLong Gold Anti-Fade Mountant (Thermo Fisher). Microscopy images were acquired at room temperature on a Leica TCS SP8 Confocal Microscope (Leica Microsystems) with a high-contrast Plan Apochromat 63×/1.30 Glyc CORR CS objective (Leica Microsystems) and an ORCA-Flash4.0 digital camera (Hamamatsu Photonics). Confocal images were acquired with Leica Application Suite X (LASX) 3.5.7 software (Leica Microsystems), followed by deconvolution using Huygens Essentials software (Scientific Volume Imaging). Representative confocal images were individually adjusted for brightness and contrast in Photoshop CC 2022 (Adobe).

### Statistical analysis

GraphPad Prism 10 was used to perform statistical analysis. One-way ANOVA was used to determine the P values of raw data with Tukey’s multiple comparison test. P values are reported in figure legends. ns = p > 0.05, * = p ≤ 0.05, ** = p ≤ 0.01, *** = p ≤ 0.001, **** = p ≤ 0.0001.

### Molecular Dynamics Simulations

All MD simulations were carried out starting from the AlphaFold2 predicted model of ABHD17A (Uniprot accession number: Q96GS6) and a simple PM-like model membrane constituted of 40%POPC-30%POPS-30%CHOL. CG-MD simulations were carried out using GROMACS package (version 2021.2) (Abraham *et al*., 2015) and employing the MARTINI3 forcefield (Souza *et al*., 2021). The initial system set up, consisting of ABHD17A positioned ∼5nm above a PM-like membrane, solvated with a 0.15 NaCl solution, was built using the insane.py script (Wassenaar *et al*., 2015) and subsequently energy-minimized using the steepest descent algorithm. After a 10 ns equilibration conducted in the NPT ensemble, four replicas of 4 μs each were run with a timestep pf 20 fs. Using semi-isotropic pressure coupling, the pressure was kept at 1 bar using the Parrinello–Rahman barostat (Parrinello & Rahman, 1981), applied every 12.0 ps with a compressibility of 3 × 10^−4^ bar^−1^. Temperature was maintained constant at 310 K employing the V-rescale thermostat (Bussi *et al*., 2007).

AA-MD simulations were started using a representative snapshot of CG simulations, where ABHD17A was stably anchored to the membrane through both N-terminus and the structurally adjacent loop. The system was first back-mapped from MARTINI3 CG model to the CHARMM36 AA model (Klauda *et al*., 2010), employing the CG2AT2 tool (Vickery & Stansfeld, 2021), and later palmitoylated at 4 sites (CYS10-11-14-15) using CHARMM-GUI tool PDB reader & manipulator (Jo *et al*., 2008; Jo *et al*., 2009). Systems containing WT and mutants were equilibrated following the classical CHARMM-GUI 6 step-protocol and then simulated using GROMACS software and the CHARMM36m forcefield. Productions were repeated three times for 500 ns each with a timestep of 2 fs in the NPT ensemble. The Nosé-Hoover thermostat (Evans & Holian, 1985) was used to keep the temperature at 310K and the Parrinello-Rahman barostat (Parrinello & Rahman, 1981) was employed, applying a semi-isotropic pressure coupling, to maintain the pressure at 1 bar and a compressibility of 4.5 x 10^-5^ bar^-1^ every 5 ps. The Particle Mesh Ewald (PME) (Darden *et al*., 1993; Essmann *et al*., 1995) algorithm was utilized for long-range electrostatic interactions, with a cutoff of 1.2 nm. Lennard-Jones (LJ) interactions were truncated at 1.2 nm. Bond constraints were treated using Linear Constraint Solver (LINCS) algorithm (Hess *et al*., 1997).

To investigate the role of flexibility of the N-terminus and the loop, AA simulations of WT-ABHD17A in solution were performed. The protein was put in a box of water with 0.15M of NaCl and minimized using steepest descent algorithm. 6-step CHARMM-GUI equilibration and 1.5μs NPT production were performed utilizing the same temperature conditions as described before for AA-simulations. The pressure was kept at 1 bar through the use of an isotropic barostat (Parrinello & Rahman, 1981) with a compressibility of 4.5 x 10^-5^ bar^-1^ using and τP=5ps).

Protein-membrane contact analysis was conducted on CG systems using ProLint (Sejdiu & Tieleman, 2021), while the occupancy map was generated using GROMACS module *gmx select*. Insertion depth for AA systems was evaluated as in Rogers & Geissler (Rogers & Geissler, 2021). Cavity pocket detection was performed utilizing the Fpocket software (Le Guilloux *et al*., 2009), giving as a reference structure for the pocket identification the AF2 model of ABHD17A. Palmitic acid was docked onto the ABHD17A AF2 model using the SwissDock webserver (www.swissdock.ch/) using default parameters (Grosdidier *et al*., 2011a; Grosdidier *et al*., 2011b). To investigate the flexibility of N-terminus and loop regions of ABHD17A WT and mutants in solution *vs.* in membrane, the distance between the center of mass of the 2 regions was measured using the *gmx pairdist*, and the results were normalized and shown as probability histogram. All the molecular images were rendered using Visual Molecular Dynamics (VMD) software (Humphrey *et al*., 1996). Plots were generated using the python module *matplotlib* (Hunter, 2007).

## Supplemental material

Figure S1 (complementary to Figure 1) shows that ABHD17A requires specific N-terminal cysteine residues for activity, acylation and plasma membrane localization. Figure S2 (complementary to Figure 5) shows the effect of mutants on the conformation, activity and binding pocket accessibility of ABHD17A. Table S1 is a list of the plasmids used in this study. Table S2 is a list of the primers used in this study.

## Acknowledgements

We thank Dr. Nevin Lambert (Augusta University, Georgia, USA) for the Venus-tagged organelle markers, Rluc8 plasmids (Lan *et al*, 2012) and guidance in quantification and statistical analysis of BRET. We also thank Dr. Alexis Shih for helping to develop the IDFP-cABPP assay. This work was supported by funding from the Canada Foundation for Innovation (Leading Edge Fund 30636, to EC), and the Canadian Institutes of Health Research (grant 162184 to EC). This work was supported by funding from the European Research Council under the European Union’s Horizon 2020 research and innovation program (grant agreement no. 803952 to SV). This work was supported by grants from the Swiss National Supercomputing Centre under projects ID s1132, s1176 and s1221.

## Author contributions

S. Holme and L. Conibear performed the *in vitro* research and analyzed the respective data. J. Sapia and S. Vanni performed the *in silico* research and analyzed the respective data. S. Holme, J. Sapia, S. Vanni and L. Conibear wrote the manuscript.

## Abbreviations

ABHD17: alpha/beta hydrolase domain-containing 17
AF2: AlphaFold2
APT: acyl protein thioesterase
BRET: bioluminescence resonance energy transfer
cABPP: competitive activity-based protein profiling
FP: fluorophosphonate
IDFP: isopropyl dodecylfluorophosphonate
MD: molecular dynamics
mSH: metabolic serine hydrolase
PAT: palmitoyl-acyl transferase
PSD-95: post synaptic density-95
STDEV: standard deviation
17-ODYA: 17-octadecynoic acid

---

**Figure S1.**
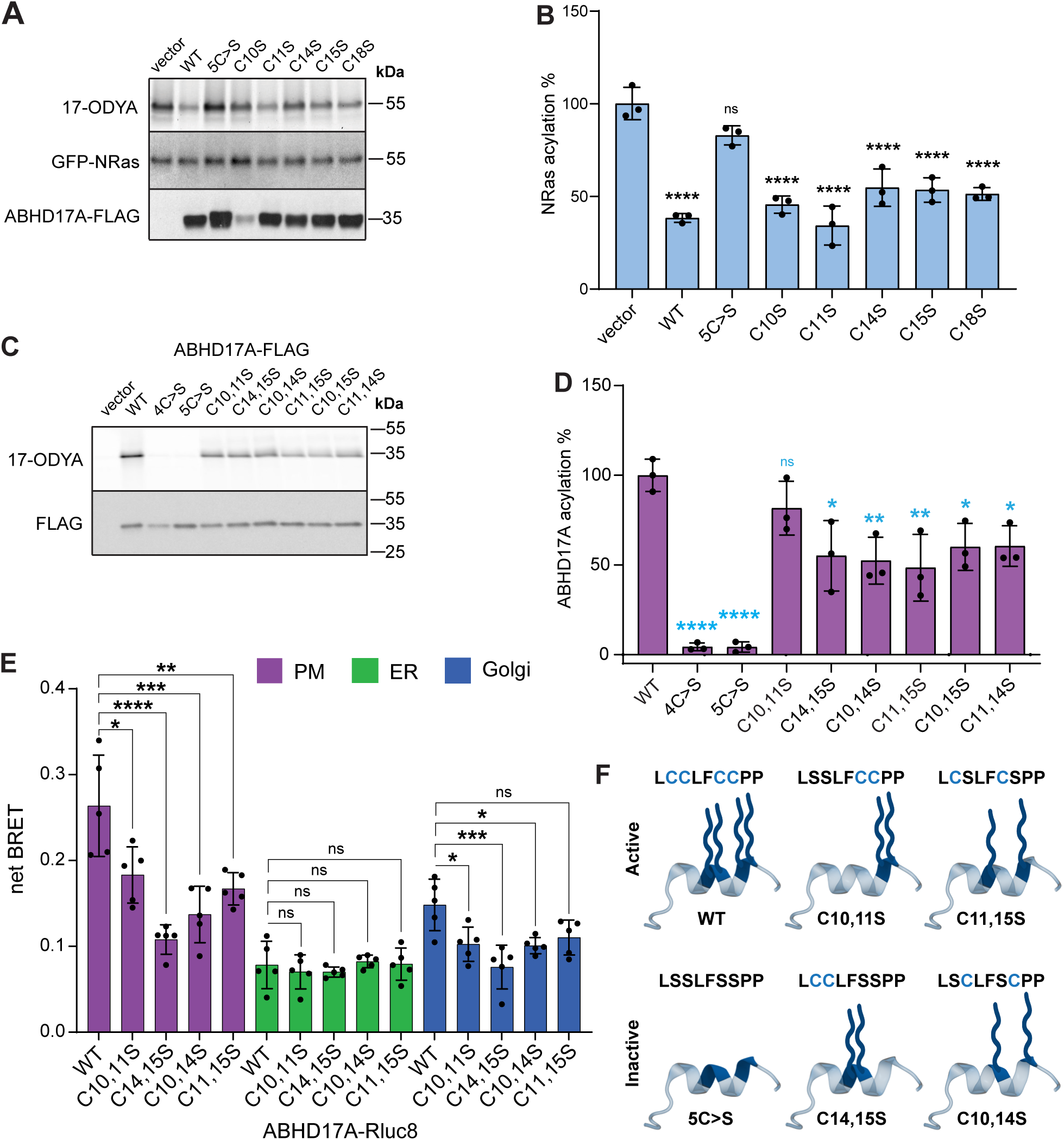
Specific cysteine residues are required for activity and plasma membrane localization. **(A)** Activity of ABHD17A is not affected by individual mutation of N-terminal cysteine residues to serine. Acylation of GFP-NRas as detected by click-labeling with 17-ODYA (upper panel). Levels of immunoprecipitated NRas (middle panel), or total ABHD17A (lower panel), were detected by western blot using anti GFP or FLAG antibody, respectively. **(B)** Quantification of GFP-NRas acylation in *(A).* One-way ANOVA with Tukey’s multiple comparison test; n=3, ns = p > 0.05, **** = p ≤ 0.0001. Statistical analysis show comparison to vector. Error bars indicate STDEV. **(C)** ABHD17A-FLAG acylation was reduced to approximately 50% for most double cysteine mutants. Acylation of ABHD17A-FLAG was detected by click-labeling with 17-ODYA (upper panel). Levels of immunoprecipitated ABHD17A were detected by western blot using anti FLAG antibody (lower panel). **(D)** Quantification ABHD17A-FLAG acylation in *(B)*. One-way ANOVA with Tukey’s multiple comparison test; n=3, ns = p > 0.05, * = p ≤ 0.05, **** = p ≤ 0.0001. Statistical analysis shows comparison to wildtype ABHD17A. Error bars indicate STDEV. **(E)** Representative double cysteine mutant constructs show decreased plasma membrane association in BRET analysis. One-way ANOVA with Tukey’s multiple comparison test; n=5, ns = p > 0.05, * = p ≤ 0.05, ** = p ≤ 0.01, *** = p ≤ 0.001, **** = p ≤ 0.0001. Error bars indicate STDEV. **(F)** Schematic of the predicted N-terminal helix of ABHD17A cysteine mutants showing acylation patterns of active and inactive constructs. Acyl groups are depicted with blue wavy lines.

**Figure S2.**
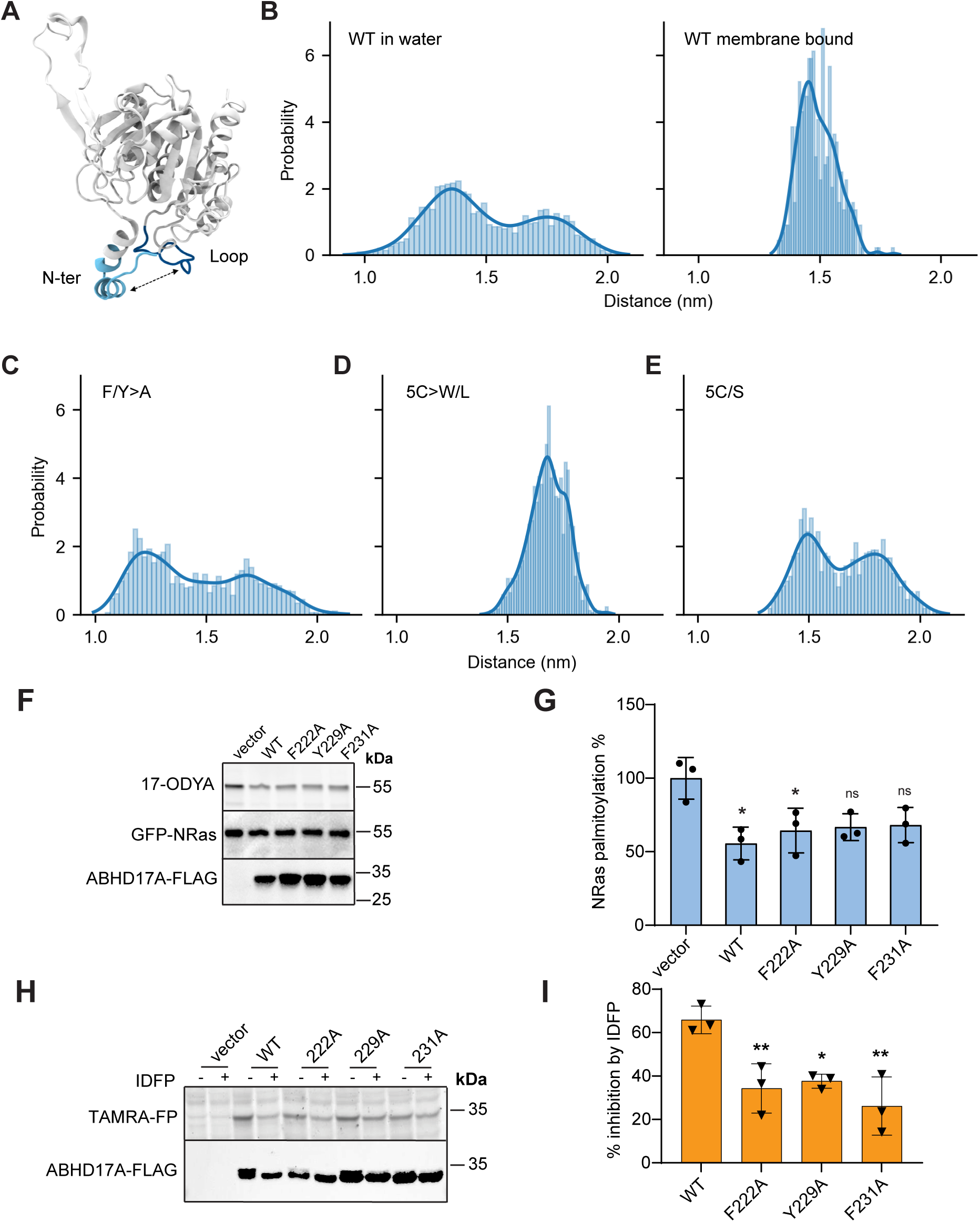
Effect of mutations in N-terminus or loop on ABHD17A conformation. **(A)** The N-terminus and nearby loop, which are responsible for the binding of ABHD17A to the membrane, are localized at the entrance of the hypothesized substrate binding pocket. **(B)** The probability distributions of the distance between the N-terminus and loop, in water (left) vs. bound to the membrane (right), indicate that the opening of the cavity displays conformational flexibility, which could influence substrate entry/exit. **(C-E)** The probability distributions of the distance between the N-terminus and loop of ABHD17A mutants. When the hydrophobic loop residues are mutated to alanine **(C)** or N-terminal cysteine residues are mutated to serine **(E)**, the N-terminus and loop regions are more dynamic, likely due to decreased membrane binding. When the hydrophobicity of the N-terminus is restored in the 5C>W/L mutant **(D)**, the enzyme displays a smaller range of conformations, which may be optimal for substrate binding. **(F)** Individual mutation of hydrophobic loop residues to alanine has little effect on ABHD17A activity. GFP-NRas acylation was detected by 17-ODYA click-labeling. Levels of immunoprecipitated NRas (middle panel), or total ABHD17A (lower panel), were detected by western blot using anti GFP or FLAG antibody, respectively **(G)** Quantification of NRas acylation in *(A)*. One-way ANOVA with Tukey’s multiple comparison test; n=3, ns = p > 0.05, *= p ≤ 0.05. Statistical analysis depicts comparison to vector. Error bars indicate STDEV. **(H)** Individual mutation of the hydrophobic loop residues to alanine has a small effect on substrate binding in cABPP analysis. **(I)** Quantification of *(C)* depicted by percentage of IDFP inhibition. One-way ANOVA with Tukey’s multiple comparison test; n=3, * = p ≤ 0.05, ** = p ≤ 0.01. Statistical analysis shows comparison to wildtype ABHD17A. Error bars indicate STDEV.

## Notes

### Competing Interest Statement

The authors have declared no competing interest.

